# Regulation of BACH1 by hemin improves cardiac function in a mouse model of myocardial infarction

**DOI:** 10.1101/2023.10.03.560676

**Authors:** Valeria Alvino, Rajesh Katare, Annabell Fricker, Elisa Avolio, Massimo Caputo, Paolo Madeddu, Sadie Slater

## Abstract

**Aims:** The BTB and CNC homology 1 (BACH1) transcription factor is a repressor of heme oxygenase-1 (HMOX1), a pivotal enzyme involved in antioxidant response and iron recycling. Here we investigated whether pharmacological modulation of the BACH1 by hemin impacts on antioxidant responses and reparative angiogenesis in a mouse model of myocardial infarction (MI).

**Methods and results:** *In vitro* studies on vascular cells showed hemin treatment downregulates BACH1 gene and protein expression and upregulates HMOX1. This axis was confirmed to be modulated in the murine infarcted heart, with BACH1 being upregulated, and HMOX1 downregulated compared to sham. Treatment with hemin every 3 days for 28 days post-MI significantly decreased BACH1 and increased HMOX1 protein expression, though no decrease in oxidative stress markers was detected. Hemin treated mice showed increases in both capillary and arteriole density, and reduced iron accumulation compared with controls. Furthermore, echocardiology measurements showed hemin treatment induced significant improvements in left ventricular wall thickness, and cardiac function as indicated by increased ejection fraction, fractional shortening, and stroke volume measurements.

**Conclusion:** Hemin has therapeutic potential to improve revascularisation and cardiac function in the heart post-MI.

## Introduction

Effective recovery from myocardial infarction (MI) depends on the balance between ongoing ischemic and metabolic damage and reparative processes. Therefore, there is significant research interest focusing on master signalling pathways, like the Bric à brac and cap’n’collar homology 1/nuclear factor erythroid 2 (NFE2)-related factor 2 (BACH1/NRF2) axis, which exert key roles in the regulation of both stress and repair.

BACH1, a transcriptional repressor, acts in opposition to the transcriptional activator NRF2, with both competing for binding to similar antioxidant response elements (ARE) on the promoter regions of target genes (e.g., HMOX1, NQO1, SOD3)^1-4^. Under normal cellular conditions, BACH1 binds to AREs of target antioxidant genes, repressing their expression, whilst NRF2 is located in the cytoplasm bound to its suppressor Kelch-like ECH-associated protein 1 (KEAP1). In this complex, NRF2 is ubiquitinated and targeted for proteasome degradation. However, in response to oxidative stress, NRF2 translocates to the nucleus, where it competes with BACH1 for binding to the ARE on the promoter region of target antioxidant genes inducing expression^1, 4, 5^.

The action of BACH1 can also be modified by heme. When cellular concentrations of heme are low BACH1 represses gene expression, however in the presence of higher concentrations of heme the BACH1 repression of genes is inactivated by BACH1 binding to heme rather than the ARE regions of target genes, therefore removing the repression and allowing their transcription^6, 7^. Alongside the well-characterised role of the BACH1/NRF2 axis in the antioxidant response, more recently it has been suggested this axis could also regulate angiogenesis. There has long been a recognised association between oxidative stress and angiogenesis. Whilst low levels or transient increases in ROS have been linked to the activation of signalling pathways instrumental to vessel growth and regeneration, high or chronic levels of ROS are detrimental to this process^8^.

We have previously demonstrated the BACH1/NRF2 signalling pathway plays a key role in the modulation of the pro-angiogenic factor angiopoietin-1 (ANGPT1) and angiogenic network formation by adventitial pericytes^9^. BACH1 has also been shown to modify ANGPT1^10^ and vascular endothelial growth factor C (VEGFC)^11^ expression *in vivo*. Furthermore BACH1 has been implicated as a modulator of reparative angiogenesis in mouse models of hindlimb ischaemia^10, 12, 13^ and a zebrafish model of developmental angiogenesis^14^ via reduced IL-8 signalling. Studies of BACH1 expression support the rationale of a BACH1 inhibitory strategy for improving cardiovascular outcomes in at risk populations. BACH1 is reportedly upregulated in heart, liver, cartilage and lungs of aging mice^15^ as well as in human bronchial epithelial cells from older adults compared to young adult donors^16^. Additionally, analysis of microarray data from healthy and acute MI patients has shown a 3.5-fold increase in BACH1 expression in MI patients circulating endothelial cells^13^.

Hemin (the active ingredient of an approved drug for treating blood disorders, such as porphyria) is an oxidised version of heme^17^. Lee *et al*^18^ recently demonstrated that the use of hemin lead to the modulation of BACH1 expression *in vitro* in breast cancer cells. This group also generated a heme-resistant mouse BACH1 mutant mouse model, which further validated that hemin acted specifically through modification of BACH1. Here we report new data showing hemin exerts a significant therapeutic action in a murine model of MI.

## Methods

### Study approval

Studies using human cells were covered by the North Somerset and South Bristol Research Ethics Committee approval (11/SW/0122) and complied with the principles stated in the “Declaration of Helsinki”. Cardiac pericytes (CP) were isolated from cardiac leftovers from infants and children undergoing palliative repair of congenital cardiac defects as previously described^19^. Informed written consent was given by paediatric patients parents/guardians for experimental use of donated material.

Animal studies were covered by license from the University of Otago, New Zealand (AEC10/14), and complied with the EU Directive 2010/63/EU. Procedures were carried out according to the principles stated in the NIH *Guide for the Care and Use of Laboratory Animals* (National Academies Press, 2011). Termination was conducted according to humane methods outlined in the Guidance on the Operation of the Animals (Scientific Procedures) Act 1986 Home Office (2014). The report of results is in line with the ARRIVE guidelines.

### Cell culture

CP were isolated from atrial or ventricle specimens (3-5mm, <100mg) with the antigenic profile confirmed by FACS as previously described^19^. Coronary artery endothelial cells (CAEC) were purchased from Promocell. BACH1 expression was modulated by hemin dissolved in ammonium hydroxide (NH_4_OH), then added to culture media at a final concentration of 1µM for the timepoints described. Vehicle treated cells were exposed to NH_4_OH only in culture media. To induce hypoxia and starvation CPs were incubated at 2% oxygen (Multi-gas incubator, MC0-19M-PE, Panasonic) for the timepoints described in EGM-2 media (Promocell) without foetal bovine serum (FBS). Conditioned media (CM), protein and RNA were then collected for further study. All experiments were performed at passage 7 (P7).

### Cell viability

Viability of CP and CAEC exposed to hemin or vehicle was evaluated using a Viability/Cytotoxicity Assay Kit (Biotium), according to manufacturer’s guidelines. Cytoplasmic Calcein-AM identified live cells, while nuclear Ethidium Homodimer III (EthD-III) the dead cells. Cells treated with 0.1% w/v saponin for 10 min served as positive control for EthD-III staining. *Caspase* CP and CAEC apoptosis was determined after treatment with hemin or vehicle using a Caspase glo kit (Promega) according to manufacturer’s guidelines. 50µM hydrogen peroxide (H_2_O_2_) was used as a control to induce cell death.

### RNA extraction and quantitative real time analysis

Total RNA was extracted using a miRNeasy kit (Qiagen). RNA concentration and purity were assessed using a NanoDrop 2000 Spectrophotometer (ThermoFisher Scientific). For mRNA analysis RNA was reverse-transcribed using a High-Capacity RNA-to-cDNA Kit (Applied Biosystems). cDNA amplification was performed on a QuantStudio 5 (ThermoFisher Scientific) and normalised to *ACTIN (Hs01060665_m1)*. Expression of *BACH1 (Hs00230917_m1), HMOX1 (Hs01110250_m1), ANGPT1 (Hs00375822_m1), VEGFA (Hs00900055_m1) a*nd *NOQ1 (Hs00168547_m1)* (all Applied Biosystems) were measured. The mRNA expression level was determined using the 2−Δct method. Each reaction was performed in triplicate.

### Protein extraction

Left ventricular tissue or cells were lysed in RIPA buffer containing phosphatase and proteinase inhibitors (1:100, both Sigma). After incubation on ice lysates were centrifuged at 12,000*xg* for 10 min at 4°C. Protein concentration was quantified using a BCA protein assay (ThermoFisher Scientific).

#### Western blotting

Protein samples were prepared in Laemmli loading buffer, incubated for 10 minutes at 95°C, resolved on 7.5 or 12% SDS-PAGE gels and transferred onto PVDF membranes (Immobilon-P PVDF Transfer Membrane 0.45, Millipore). Membranes were blocked using 5% BSA or 5% non-fat dried milk in TBS containing 0.05% Tween 20 (Sigma-Aldrich) for 1h at 15-25°C. Primary antibodies for BACH1 (1:1000, Proteintech), HMOX1 (1:1000, Cell Signalling Technologies) and loading controls TUBULIN (1:1000, Cell Signaling Technologies) or ACTIN (1:5000, Sigma), along with corresponding species IgG secondary antibodies (1:5000, GE Healthcare, Fisher Scientific) were utilised. Membrane development was performed by an enhanced chemiluminescence-based detection method (ECL™ Prime Western Blotting Detection Reagent, GE Healthcare, Fisher Scientific) using a ChemiDoc MP system (Bio-Rad). Blot densitometry was analysed using Image J 5.1 software, normalised to Actin, Tubulin or total protein expression detected using Ponceau staining.

### ELISA

CP were incubated in EGM-2 media without FBS for 48h under normoxic or hypoxic conditions, or with hemin treatment as described above. CM was collected and secreted ANGPT1, ANGPT2 and VEGFA levels were analysed using manufacture’s ELISA protocol (Bio-Techne).

### Angiogenesis array

CP were treated with either vehicle or hemin and CM was collected as descripted above. Expression of angiogenic proteins in CM was measured using a Proteome Profiler Human Angiogenesis Array kit (ARY007, R&D systems) as per the manufacturer’s instructions. Membrane development was performed by an enhanced chemiluminescence-based detection method (ECL™ Prime Western Blotting Detection Reagent, GE Healthcare, Fisher Scientific) using a ChemiDoc MP system (Bio-Rad). Blot densitometry was analysed using Image J 5.1 software, normalised to array controls.

### Mouse model of chronic myocardial infarction (MI) and hemin treatment

Following basal echocardiography (GE Vivid E9), MI was induced in 8-weeks old C57/Bl6 male mice (Hercus) by permanent ligation of the left anterior descending coronary artery (LAD) as described previously^20, 21^. In brief, following anaesthesia (2,2,2 tribromo ethanol, 0.3gm/kg, i.p.) and artificial ventilation, the chest cavity was opened and, after careful dissection of the pericardium, LAD was located and permanently ligated using a 7-0 silk suture. After confirming for the absence of bleeding, chest cavity was closed in layers. Animals were allowed to recover for at least 4 hours before returning to the housing unit. Mice were monitored twice a day for the first 5 days post-surgery and thereon once every day. All the mice received analgesic and antibiotic from just before the surgery and for the next 3 days. For initial experiments to determine BACH1 and HMOX1 expression, on day 10 post surgery, mice were euthanised and ventricular tissue collected for Western blot analysis. For the treatment group on day 1 post surgery animals were randomized to receive vehicle or three different doses of hemin (12.5mg/kg, 25mg/kg and 50mg/kg, Sigma Aldrich) intraperitoneally every 3 days until 28 days post-MI. NH_4_OH was used as vehicle. On day 28, following echocardiography and under anaesthesia, heart were stopped in diastole using potassium chloride, and ventricular tissue was collected for Western blot and histological analysis.

### Histological analysis

Paraffin-embedded sections of mouse heart tissue were cut using a Shandon Finesse 325 manual microtome (ThermoFisher) and assessed for HMOX1 expression using an anti-mouse HMOX1 primary antibody (1:100, Abcam, ab13243). Before addition of the primary antibody the sections were subjected to antigen retrieval using citrate buffer 0.02 mol/L, pH = 6 for 30 min at +98°C. The 3,3′-Diaminobenzidine (DAB) staining method was used to identify the HMOX1 antibody in form of brown precipitates. Iron deposits were stained using an iron staining kit according to the manufacturer’s instructions (Abcam, ab150674). Nuclei were counterstained using Mayer Haematoxylin (MHS16-500ML) followed by dehydration and mounting in DPX resin for the imaging. HMOX1 immunostaining and iron deposits were visualised using light microscopy and the images were taken using ImageJ software.

To determine the production of hydroxyl radicals seven-micron thick left ventricular cryosections were probed with polyclonal 8-hydroxy-2’-deoxyguanosine (8-OHdG) antibody overnight at 4°C, followed by incubation with goat anti-rabbit secondary antibody conjugated with Alexa 568 (Thermofisher) at room temperature for 3 hours. DAPI (1:1,000 dilution, Santa Cruz Biotechnologies) was used to stain the nuclei. Images were captured as above and expressed as % of 8-OHdG/field.

To assess capillary and arteriole density, cryosections were probed with biotin-conjugated Isolectin-B4 (1:200 dilution, Vector laboratories) to detect endothelial cells (to determine the capillary density) and anti-α-smooth muscle actin conjugated with Cy3 (1:800 dilution, Sigma-Aldrich) to detect smooth muscle cells (to determine the arteriole density), respectively. DAPI (1:1,000 dilution, Santa Cruz Biotechnologies) was used to stain the nuclei. Images were captured at 200× magnification using a fluorescence microscope (Olympus). The density of capillaries was expressed as the mean number of isolectin^+^ cells/mm^2^ of cardiac tissue. The density of arterioles (<50 μm lumen size) was expressed as the mean number of αSMA^+^ isolectin^+^ cells/mm^2^ of tissue.

### Statistical analysis

All data were analysed using GraphPad Prism software (v9). Two-group analysis was performed by Student’s t-test. Comparison of multiple groups was performed by analysis of variance (ANOVA) with post-test multiple comparisons tests. Values were expressed as means ±SEM. Probability values (P) less than 0.05 were considered significant.

## Results

### Regulation of BACH1 in vitro using hemin

Dose-response studies were performed to determine the optimal dosage of hemin to treat cardiac vascular cells (CAEC and CP) with the intent to remove BACH1 gene repression without inducing detrimental cellular effects (Data not shown). From this, it was determined the optimal dosage to treat CAEC and CP with was 1µM hemin for 24h as this concentration and timepoint had no effect on cell viability (**Figure 1A&B, E&F**), did not induce cell apoptosis (**Figure 1C&D**) or affect cell morphology (**Figure 1G&H**).

**Figure 1.**
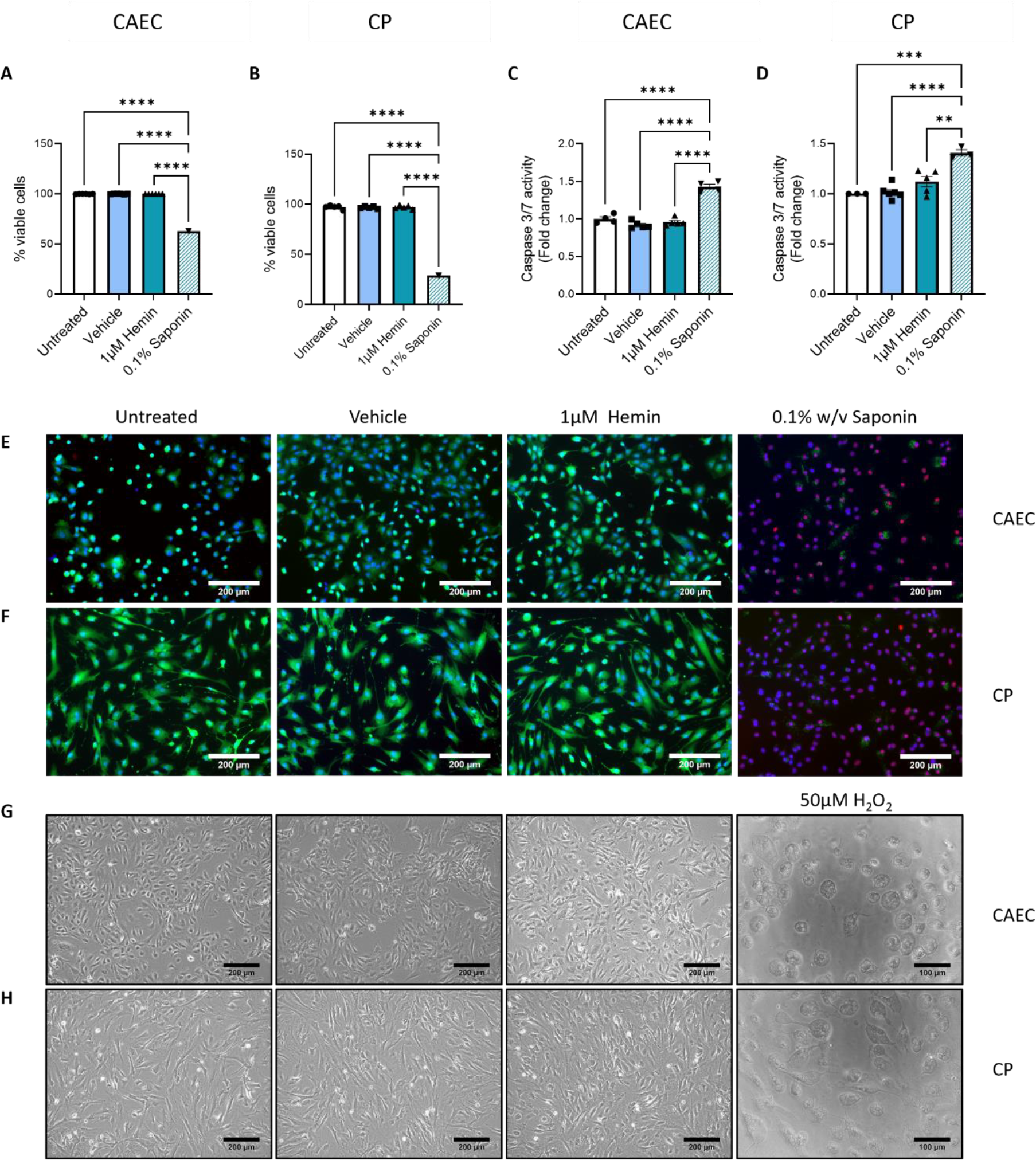
Effect on hemin treatment on cardiac vascular cells. Treatment of (**A**) CAEC or (**B**) CP with 1µM hemin for 24h had no detrimental effect on cell viability as determined by live/dead staining. (**E& F**) Representative fluorescent images. Green: viable cells, red: non-viable cells, blue: DAPI nuclear stain. 0.1% Saponin w/v was used as a control to reduce cell viability. Treatment of (**C**) CAEC or (**D**) CP with 1µM hemin did not induce cell death as measured by a caspase assay. Treatment with 50µM H_2_O_2_ was used as a control to induce cell death. (**G & H**) Representative light microscopy images of cells after 24h treatment with 1µM hemin. One way ANOVA, ± SEM, n=2-5. ***p<0.001 and ****p<0.0001 vs. untreated cells.

### Effect of BACH1 modulation on target factor expression

The effects of hemin treatment on expression of BACH1 and downstream target factors in CAEC and CP was then determined. There was no change in BACH1 gene expression in CAEC after 24h treatment with hemin, however BACH1 protein expression significantly decreased. (**Figure 2A-B&E**). Surprisingly, hemin treatment induced BACH1 gene expression in CP, however this was not reflected in protein expression which was significantly decreased (**Figure 2C-E**). For both CAEC and CP, hemin treatment increased gene and protein expression of HMOX1 (**Figure 2E-I**). Likewise, an increase in gene expression of NQO1 (another oxidative stress responsive enzyme regulated by BACH1) was also noted in CP (**Figure 2J&K**). These data show for the first-time the regulation of oxidative stress responsive enzymes HMOX1 and NQO1 by BACH1 in cardiac vascular cells.

**Figure 2.**
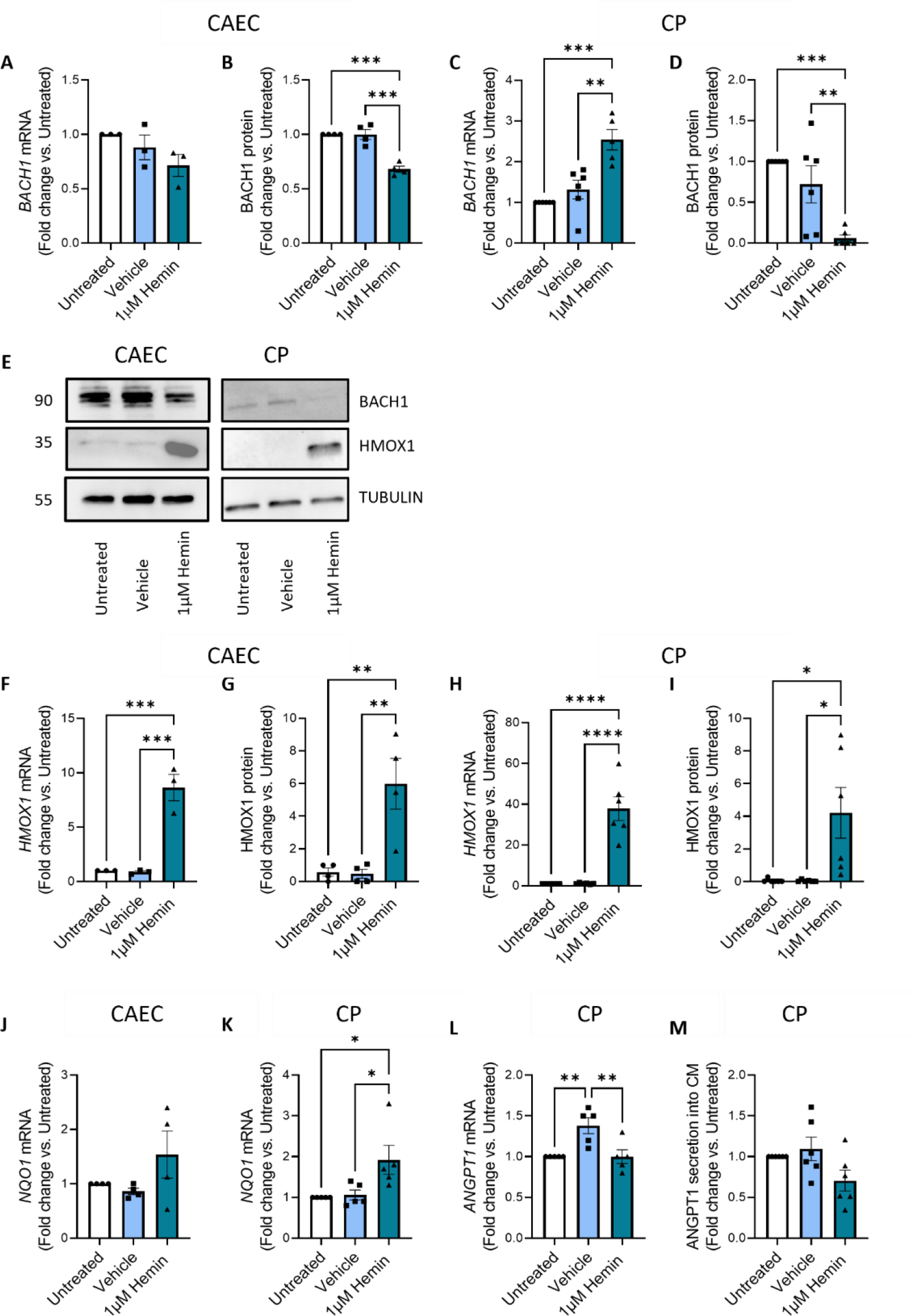
Effect of hemin treatment on BACH1 targets in cardiac vascular cells. Treatment of CAEC with hemin had no effect on BACH1 **(A)** gene expression however **(B)** protein expression significantly decreased. In CP BACH1 **(C)** gene expression significantly increased, whilst BACH1 **(D)** protein expression significantly decreased. **(E)** Representative Western blots. HMOX1 **(F & H)** gene and **(G & I)** protein expression significantly increased in both CAEC and CP respectively. NQO1 gene expression **(J & K)** only increased significantly in CP. In CP there was no change in **(L)** ANGPT1 gene expression or **(M)** ANGPT1 secretion into CM. One way ANOVA, ± SEM, n=2-5. *P<0.05, **p<0.01, ***p<0.001 and ****p<0.0001 vs. untreated cells.

In CP, there was no change in ANGPT1 gene expression or amount of ANGPT1 secreted into the conditioned media (Figure 2L-M). To check whether the effect on ANGPT1 expression was time dependent, we increased the treatment time with hemin to 48h. At this timepoint, the CP maintained the decrease in BACH1 and increase in HMOX1 protein expression and induced a significant decrease in secretion of ANGPT1 into the conditioned media (**Supplemental Figure 1**). This is in contrast to our previous data using siRNA to silence BACH1, where an increase in ANGPT1 was observed^9^. We then used siRNA to silence BACH1 in CP with the aim to check the same targets were regulated. siRNA silencing of BACH1 in CP caused a significant decrease in BACH1 gene and protein expression (**Supplemental figure 2A-B&E**), whilst HMOX1 showed a significant increase in both gene and protein expression (**Supplemental figure 2C-E**). These data confirm depletion of BACH1 by siRNA in CP gave the same results as observed when CP were treated with hemin. Additionally, there was a significant increase in ANGPT1 gene expression when CP BACH1 was silenced by siRNA although no significant change in ANGPT1 secretion into conditioned media was observed (**Supplemental figure 2F&G**). These data suggest BACH1 regulates ANGPT1 gene expression, however the repression/expression of ANGPT1 is dependent on the methodology and duration of treatment to alter BACH1 expression.

We then utilised a proteome array to determine if hemin alters angiogenic factor expression in CP. Fifty-five angiogenic factors were determined in response to treatment with conditioned media from CP with and without hemin supplementation. (**Figure 3A&B**). At this timepoint and concentration, no significant change was observed between hemin and control. Additional experiments determined there was no change in either VEGF or ANGPT2 gene expression or secretion into the CM when CAEC were treated with hemin (data not shown). Alongside this, we determined the effect of hemin on vascular network formation *in vitro*. Using a Matrigel assay and co-culture of CAEC and CP with and without hemin treatment, we found no significant change in vascular network formation when hemin treated cells were compared to untreated cells or cells treated with vehicle only. (**Supplemental figure 3A-D**). Altogether these findings suggest BACH1 modulation with hemin over 24-48h *in vitro* has no effect on expression of angiogenic factors by cardiac vascular cells or vascular network formation.

**Figure 3.**
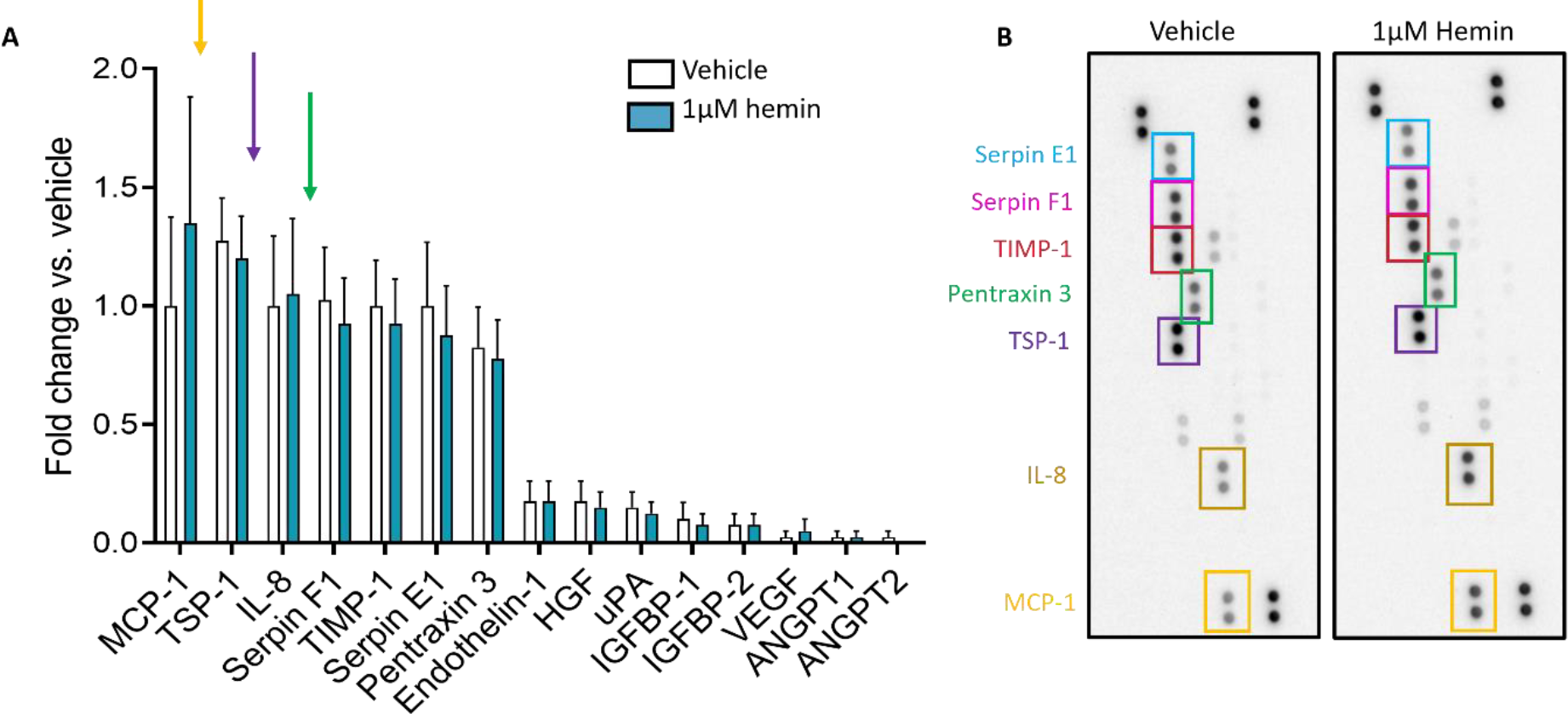
Expression of secreted angiogenic factors by CPs. **(A)** Array data showing expression of angiogenic factors by CP in response to vehicle or hemin treatment. No significant changes were observed. **(B)** Representative array images. One way ANOVA, ± SEM, n=4.

### Effect of hypoxia and starvation on CP

In this experiment, CP were exposed to hypoxia and starvation for 48h, then expression of BACH1 and HMOX1 was determined. No change in BACH1 gene expression was observed when CP were exposed to hypoxia and starvation compared to CP incubated under normoxic conditions, however BACH1 protein expression significantly increased (**Figure 4A-B&D**). A significant decrease in HMOX1 gene expression was observed with hypoxia and starvation, whilst no expression of HMOX1 protein was detected under either condition. This is to be expected as HMOX1 expression is repressed by the presence of BACH1. This data suggests that, under hypoxic conditions, the antioxidant response is impaired (**Figure 4C&D**). Interestingly, the increase in BACH1 expression induced by hypoxia and starvation also correlated with significantly decreased expression of the ANGPT1 gene, however, at this time point, ANGPT1 protein secretion into the conditioned media did not change (**Figure 4E&F**). This data suggests exposure of CP to hypoxia and starvation could impact on ANGPT1 secretion.

**Figure 4.**
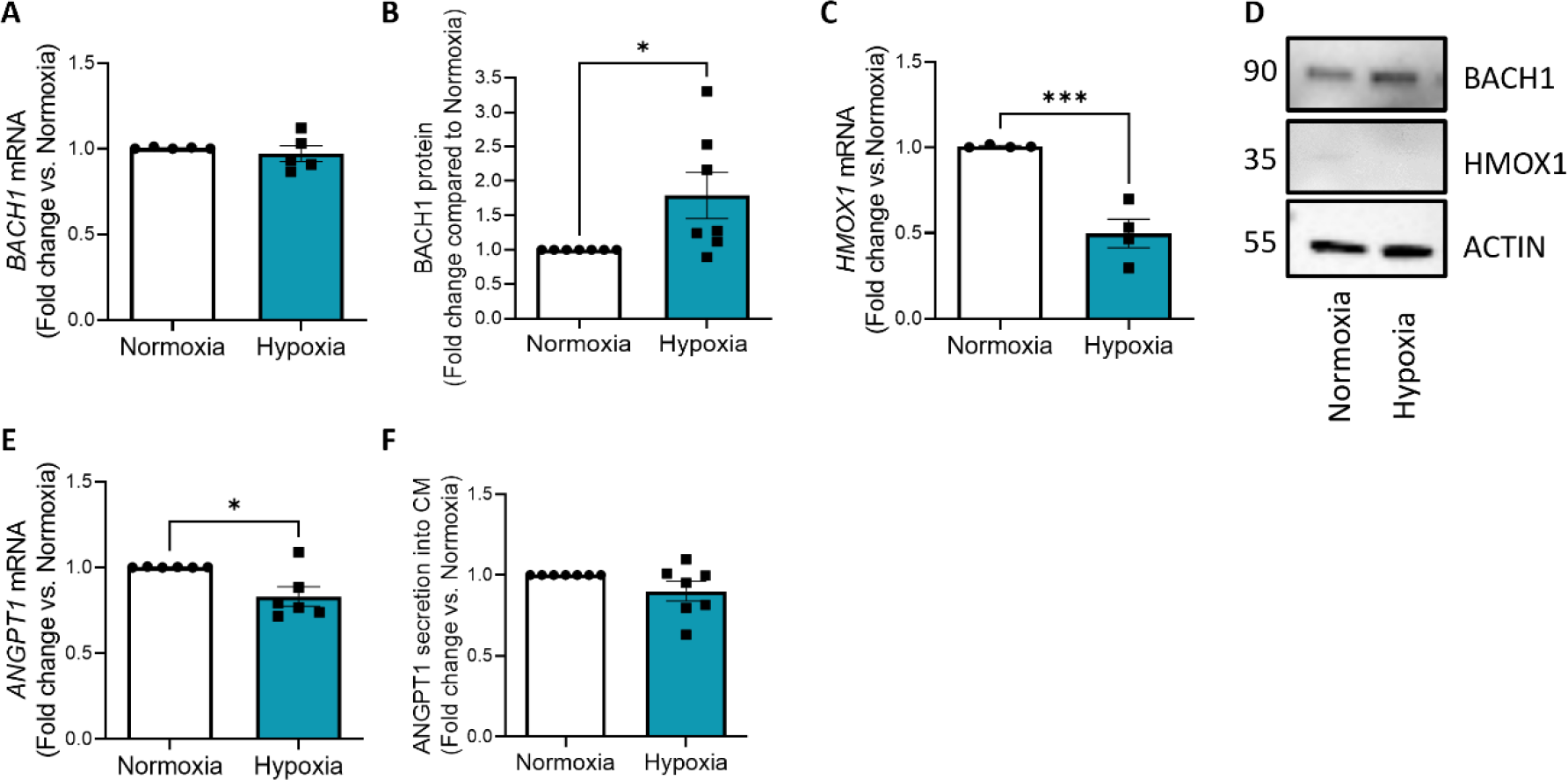
Effect of hypoxia and starvation on CP. There was no significant change in BACH1 **(A)** gene expression in response to 48h hypoxia compared to normoxic controls however **(B)** protein expression significantly increased. There was also a significant decrease in **(C)** HMOX1 gene expression. No HMOX1 protein was detectable under either condition. **(D)** Representative Western blots. ANGPT1 **(E)** gene expression was found to be significantly decreased in response to hypoxia, however **(F)** ANGPT1 secretion into the conditioned media did not change. Unpaired Student’s t-tests, n=5-6, values ± SEM. *P<0.05 and ***p<0.001 vs. normoxia.

### BACH1 and HMOX1 expression in mice after MI

We next determined protein expression of BACH1 and its downstream target HMOX1 in mouse hearts 3- and 10-days post MI. Compared to sham mice, BACH1 expression significantly increased at both timepoints. Interestingly, whilst still raised compared to sham, there was a significant decrease in BACH1 expression when day 10 post MI was compared to day 3, suggesting this is a transient effect (**Figure 5A&C**). In tandem with this, HMOX1 expression was significantly decreased at both timepoints compared to sham mice (**Figure 5B&C**), therefore suggesting in the initial period after MI the antioxidant response is suboptimal and that a rebalancing of BACH1 and HMOX1 expression may improve the antioxidant response. Deposition of iron is a recognised complication of cardiac reperfusion after MI, causing a peculiar form of cell death (ferroptosis) through iron-dependent lipid peroxidation^22^. We measured iron accumulation in histological sections taken from sham animals and 3 areas of the heart after MI (remote area - RA, peripheral ischaemic area – PIA, ischaemic area-IA). We observed a significant increase in iron accumulation in cells post MI in the infarcted area compared to sham, thus suggesting that permanent ischemia without reperfusion can also cause myocardial iron deposition (**Supplemental figure 4A&B**).

**Figure 5.**
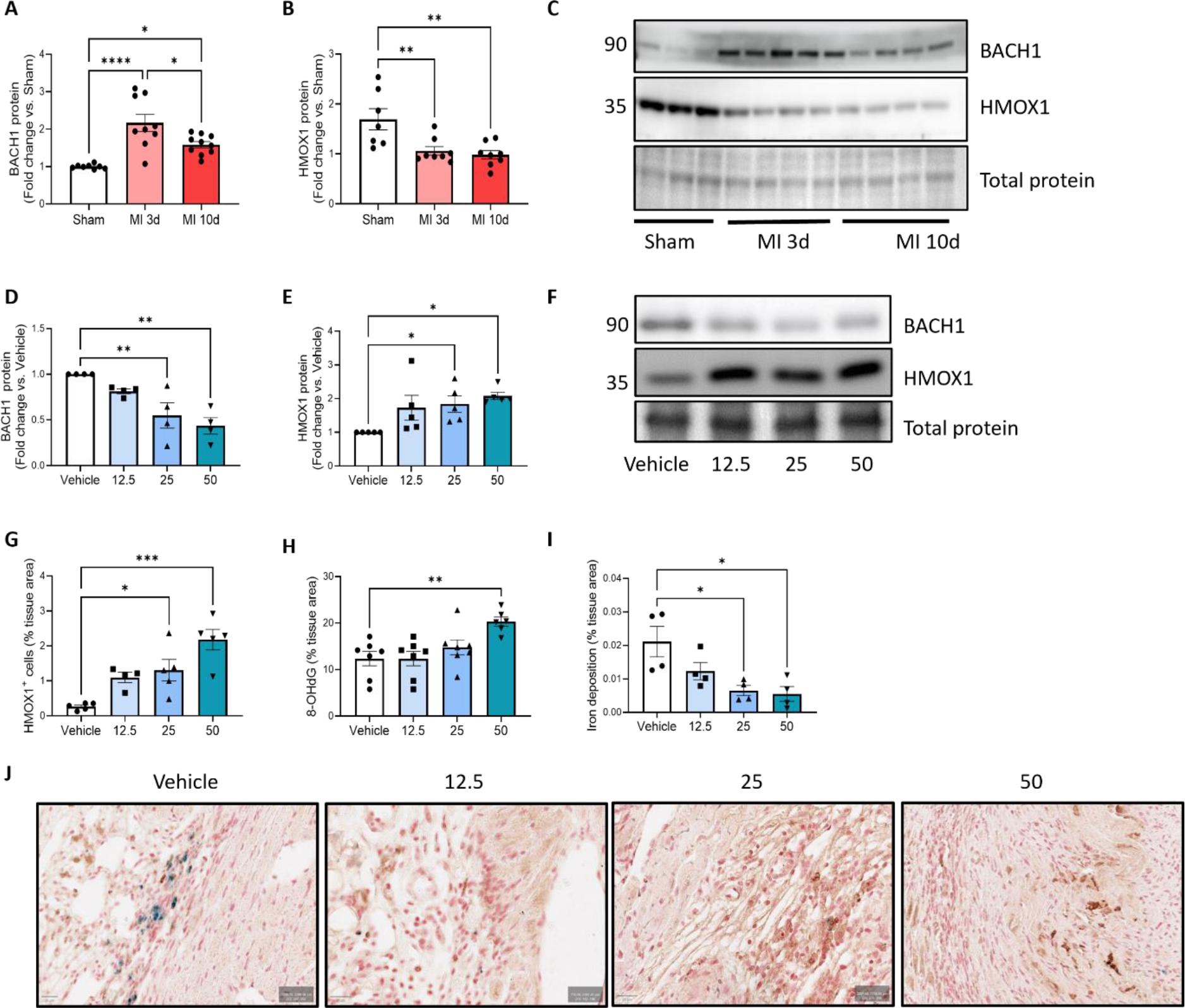
Expression of BACH1 and HMOX1 in mouse heart tissue after MI. **(A)** BACH1 protein expression significantly increased in mouse heart tissue at both day 3 and 10 post MI, whilst **(B)** HMOX1 expression significantly decreased at both timepoints. **(C)** Representative Western blots. **(D)** BACH1 protein expression significantly decreased in mouse heart tissue when mice were treated with either 25 or 50mg/kg hemin. Meanwhile HMOX1 expression significantly increased with both dosages **(E)** protein data, **(F)** representative Western blots. Staining of heart tissue sections show **(G)** HMOX1 expression significantly increases in hemin, whilst **(H)** Iron expression significantly decreases. **(I)** A significant increase in expression of oxidative stress marker 8-OHdG was observed in animals treated with the highest concentration of hemin. **(J)** Representative images of tissue staining. Brown: HMOX1, Blue: iron, purple: haematoxylin nuclear stain. One way ANOVA, ± SEM, n=7-8. *P<0.05, **p<0.01 and ****p<0.0001 vs. sham/vehicle.

### Effect of hemin treatment in mice after MI

We then determined the effects of hemin treatment on BACH1 and HMOX1 expression in the heart after MI. Mice were treated with vehicle or 3 dosages of hemin (12.5, 25 and 50mg/kg)^18^ every 3 days until 28 days post MI. BACH1 and HMOX1 expression was then determined by Western blotting. Whilst there was no significant effect with the lowest dose, treatment of mice with 25 or 50mg/kg hemin after MI resulted in a significant decrease in BACH1 and a significant increase in HMOX1 protein expression compared to vehicle, thus reversing the increase in BACH1 and repression of HMOX1 expression observed after MI (**Figure 5D-F**). Staining of tissues sections for HMOX1 also showed increased expression with hemin treatment (**Figure 5G&J**). We next determined the amount of reactive oxygen species in the heart tissues by staining for 8-OHdG. Tissues from animals treated with 12.5 and 25mg/kg hemin showed similar levels of 8-OHdG expression, whilst surprisingly, an increase in 8-OHdG expression was observed in hearts treated with 50mg/kg hemin (**Figure 5H**). Having observed a significant increase in iron accumulation in the heart after MI, we determined iron accumulation after treatment with hemin. Treatment with the higher dosages of hemin correlated with significantly less iron accumulation in the heart tissue (**Figure 5I&J**).

To determine the effect of hemin treatment on angiogenesis after MI, we quantified blood vessel density in the remote and peri-infarct regions of the heart by histological staining of ventricular sections for isolectin-B4 and anti-α-smooth muscle actin. Hemin treatment increased capillary density in peri-infarct zone with Tukey’s multiple comparisons test identifying all dosages of hemin as significantly different from vehicle (**Figure 6A&C**). Likewise, hemin treatment increased the arteriole density of the peri-infarct zone, with 25mg/kg being the lowest effective dosage and 50mg/kg inducing a further incremental effect. A multiple comparison test did not detect a different between these two dosages. (**Figure 6B&C**). Hemin treatment did not alter the capillary or arteriole density in the remote zone (data not shown). Together these findings indicate treatment with 25mg/kg hemin post MI is sufficient to induce a significant improvement in reparative angiogenesis in the infarcted region of the heart.

**Figure 6.**
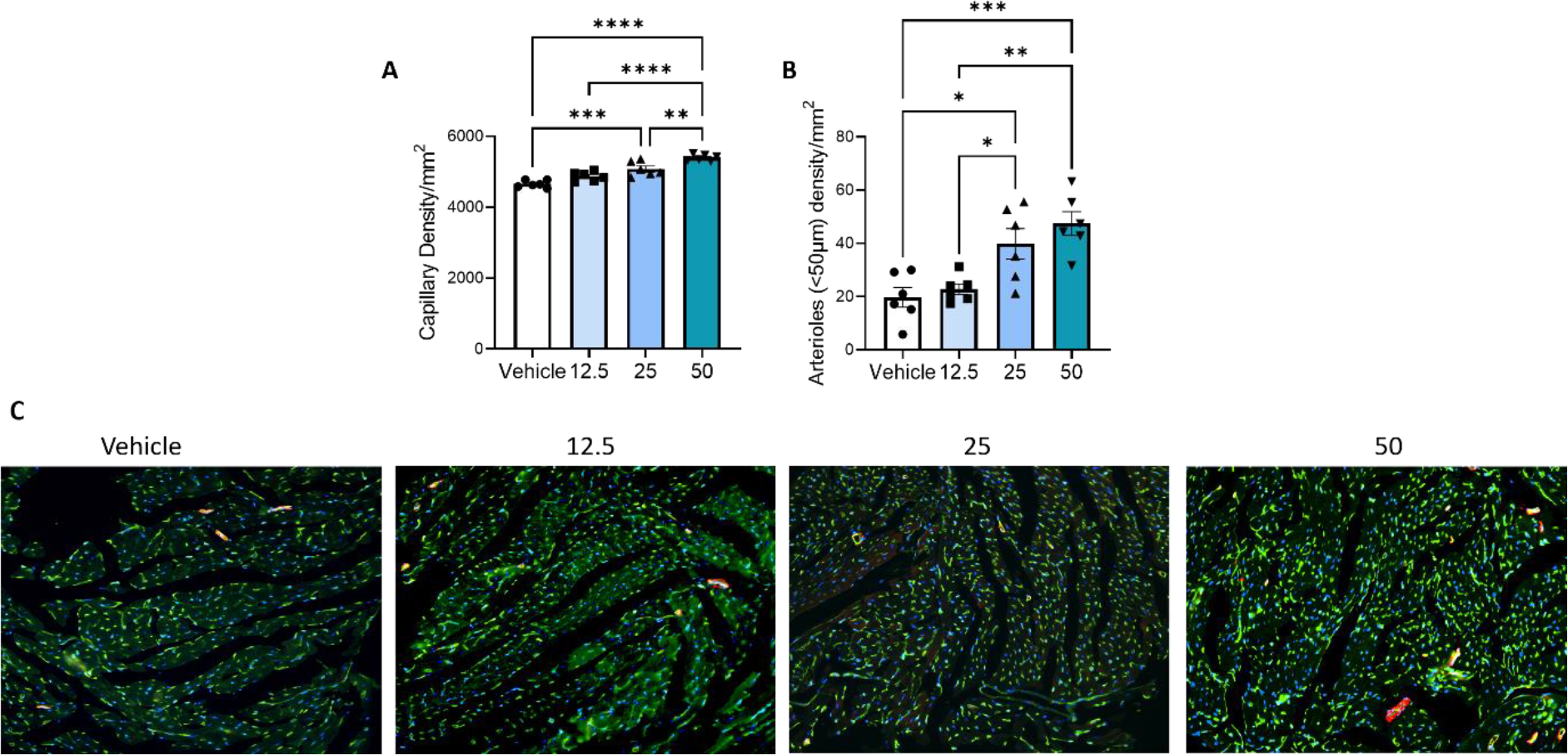
Effect of hemin treatment on capillary/arteriole density in mouse MI model. **(A)** Capillary density significantly increased with hemin treatment in the peri-infarct region, as did **(B)** arteriole density. **(C)** Representative images. Green: Isolectin B4, red: α smooth muscle actin, blue: DAPI nuclear stain. One way ANOVA, ± SEM, n=6. **p<0.01 and ****p<0.0001 vs. vehicle.

The reported anatomical effects of hemin were associated with improvements in volumetric and contractility indexes, as assessed by 3D echocardiography at baseline and 28 days post MI. Echocardiography showed hemin treatment had no effect on overall heart rate (**Supplemental Figure 5A**). In vehicle-treated mice, there was an expected thinning of the left ventricular end-diastolic anterior wall thickness (LVAW) corresponding to the infarct area, yet interestingly, a significant increase in the thickness of the LVAW was observed after 25 and 50mg/kg hemin treatment (**Figure 7A**). Alongside this an increase in LV mass/body weight ratio was also observed with hemin treatment (**Figure 7B**). No effect on left ventricular posterior wall end systole (LVPWs) was observed but a small but significant increase in left ventricular posterior wall end diastole (LVPWd) was seen with 50mg/kg hemin (**Figure 7C&D**). Moreover, hemin induced a remarkable improvement in volumetric and contractility indexes. Significant decreases in left ventricular internal diameter end systole (LVIDs), left ventricular internal diameter end diastole (LVIDd), end-systolic volume (ESV) and end-diastolic volume (EDV), were observed when mice were treated with either 25 or 50mg/kg hemin (**Figure 7E-H**). Likewise, ejection fraction (EF), fractional shortening (FS) and stroke volume (SV) all significantly increased in mice treated with either 25 or 50mg/kg hemin compared to vehicle (**Figure 7I-K**). We also tested if the benefits were dose related. Two-way ANOVA detected an interaction with dosage and post-hoc analysis identified 25mg/kg to be the lowest effective dosage. Survival curves suggested a lower number of deaths in the treated groups, however Kaplan-Meier analysis of the survival data did not detect a significant difference between hemin treatment and vehicle (**Supplemental Figure 5B**).

**Figure 7.**
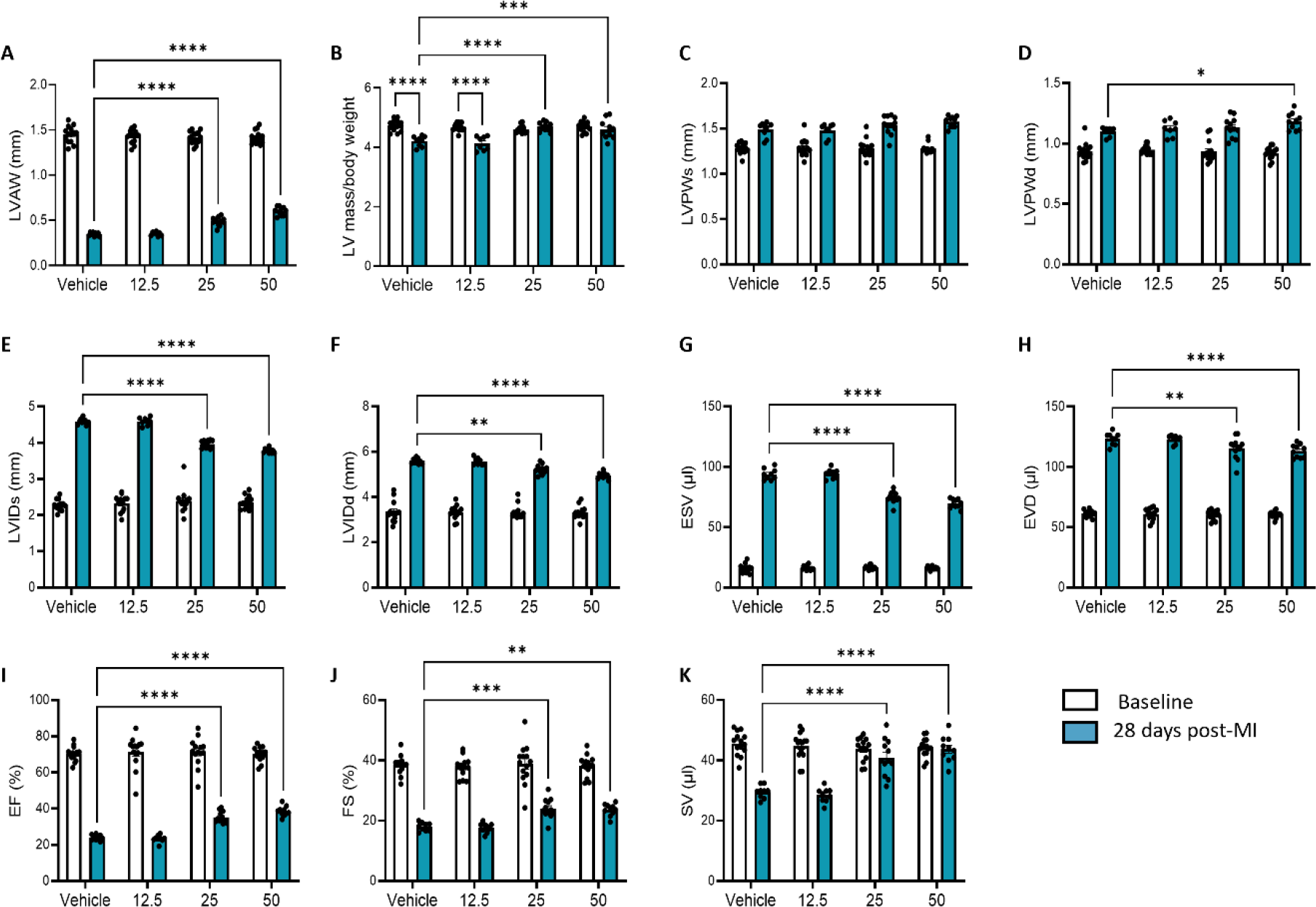
Effect of hemin treatment on cardiac functional parameters in mouse MI model. Treatment of mice with increasing dosages of hemin caused an improvement in measures of cardiac function. Treatment with 25 and 50mg/kg hemin caused significant changes in **(A)** LVAW and **(B)** LV mass/body weight ratio. Hemin treatment had no effect on **(C)** LVPWs but did significantly increase **(D)** LVPWd at the highest dosage of treatment. **(E)** LVIDs **(F)** LVIDd **(G)** ESV and **(H)** EDV all significantly decreased, whilst **(I)** EF **(J)** FS and **(K)** SV measurements all significantly improved with hemin treatment. Two-way ANOVA, ± SEM, n=14. *P<0.05, **p<0.01, ***p<0.001 and ****p<0.0001 vs. vehicle.

## Discussion

We believe this study provides a new target for inducing reparative angiogenesis and functional improvement in the heart after MI. The transcription factor BACH1 may represent a novel druggable target for the modulation of both angiogenesis and antioxidant response. Using a mouse model, we showed BACH1 expression is increased after MI and remains elevated even after 10 days, leading to repression of the antioxidant factor HMOX1 during the time the heart is most susceptible to generation of ROS through hypoxia. We also observed increased levels of cellular iron, which has also been implicated to generate oxidative stress^23^. This is in keeping with previous data showing BACH1 levels remain elevated during the first 48h post MI in the murine left ventricular myocardium^24^ and microarray data from human circulating endothelial cells demonstrating elevated BACH1 expression after MI^25^. Furthermore, other cardiovascular risk factors such as elevated glucose^26^ and aging^27^ have also been associated with increased BACH1 expression in the mouse aorta and heart respectively. For aging, this was further linked to increased endothelial cell senescence when under oxidative stress^27^. Likewise, BACH1 expression was also found to be elevated in human atherosclerotic plaques^28^. BACH1 deletion in a mouse model of atherosclerosis was associated with reduced proinflammatory cytokine expression^28^. Taken together these data suggests elevated BACH1 expression is a risk factor for a poorer outcome in response to MI for both angiogenic and antioxidant responses. Here we demonstrated BACH1 expression can be pharmacologically modulated *in vivo* using hemin, which is already approved for use therapeutically to treat porphyria, to improve cardiac outcome after MI.

BACH1 is known to have a role in regulating expression of genes involved in the antioxidant response^1, 3, 29^. In this work we found an increase in expression of HMOX1 gene and protein both *in vitro* in cardiac vascular cells and *in vivo* in the heart with BACH1 degradation, therefore we anticipated this would decrease oxidative stress markers. 8-OHdG, a product of oxidative damage to 8-hydroxy-2′-deoxyguanosine, is a well-characterised marker for assessing oxidative DNA damage. Staining heart sections surprisingly showed a significant increase in 8-OHgD in heart tissue from animals treated with the highest dosage of hemin, whilst the other dosages showed similar levels of 8-OHdG as that found in the vehicle. These results could be due to the timepoint assessed in this study. Here, we used a timepoint of 28 days to assess the effect of hemin treatment on cardiac remodelling. It is possible this time point is too late to accurately reflect the impact of hemin treatment on oxidative damage, which primarily occurs in the early days after MI^30^.

We also examined the accumulation of iron in the heart after MI. Whilst iron is essential for many biological functions, excessive accumulation is toxic to cells^22, 23^. Additionally iron accumulation is also a recognised risk factor for cardiovascular disease and a complication of myocardial reperfusion^22, 31^. We observed a significant increase in iron accumulation in the heart after MI compared to sham animals, and, interestingly, treatment with hemin significantly reduced the accumulation of iron in the heart. Further work is required to determine how hemin could regulate iron accumulation in the infarcted heart.

Alongside its role in the antioxidant response, more recently BACH1 has also been identified to modulate angiogenesis in zebrafish^14^ and mouse models of limb ischaemia^10, 12, 13^. For all models a reduction in BACH1 expression was associated with increased angiogenesis. In keeping with previously published work, our *in vivo* data showed administration of hemin induced both increased capillary and arteriole density in the peri-infarct zone of the heart. However, our *in vitro* data showed no change in network formation using a Matrigel assay, therefore the molecular mechanism for this *in vivo* increase in blood vessel density is still to be elucidated. It is challenging to replicate the milieu of factors generated in response to ischaemia and reperfusion *in vivo* in an *in vitro* model. However, *in vitro* investigations inducing hypoxia and starvation in CPs to replicate the ischaemic condition of the heart tissue post MI led to increased BACH1 and reduced HMOX1 expression, alongside decreased ANGPT1 gene expression. We hypothesise a longer exposure of cells to hypoxia/starvation would result in decreased secretion of ANGPT1 into the conditioned media. We previously demonstrated silencing of BACH1 using siRNA in adventitial pericytes increased ANGPT1 expression^9^, however we did not see this effect when degrading BACH1 expression using hemin in vascular cells *in vitro*. However, BACH1 silencing by siRNA in CPs induced an increase in ANGPT1 gene expression and a trend for an increase in ANGPT1 secretion. This anomaly in ANGPT1 regulation may be due to the mechanism used to modulate BACH1. Whilst the siRNA silences BACH1 gene expression, hemin reduces the effects of BACH1 by inducing BACH1 binding to heme rather than gene promoter regions of target genes. These data still suggest BACH1 regulation of ANGPT1 gene expression.

Furthermore, Yusoff *et al*^10^ has also demonstrated an increase in ANGPT1 expression in a BACH1-/-mouse model of hind limb ischaemia, which correlated with increased capillary density in the BACH1-/-mouse compared to wild type. Additionally, BACH1 has also been linked to regulation of other angiogenic factors. In zebrafish, BACH1 has been demonstrated to regulate angiogenesis and lymphangiogenesis via modulation of VEGFC^11^.

Finally, our *in vivo* studies further support a role for hemin as a therapy post MI due to the improvements observed in cardiac function. Our data showed hemin treatment induced increases in LVAW thickness alongside increases in LV mass, suggesting hemin induces remodelling of the scar region. Moreover, hemin treatment induced significant improvements in contractility indexes ejection fraction, fractional shortening and stroke volume, which could be at least in part attributed to the improvement in neovascularization.

## Conclusion

Our findings suggest that hemin treatment can help improve cardiac vascularization and function after MI. Whilst further work is required to fully elucidate a mechanism, the data suggests a role for BACH1. As BACH1 is a known regulator of genes involved in the angiogenic and oxidative responses upregulation of the genes involved in these processes would be beneficial in the recovery of tissue after an ischaemic insult. As mentioned previously, further work using early timepoints post MI are required to fully assess the effectiveness of hemin treatment on modulating the oxidative stress response. Furthermore, additional investigations on the effect of hemin treatment on fibrosis would also be informative to determine if these effects are due to scar remodelling or improvements in muscle mass. However, the functional histology and survival data all show a preponderance of benefit versus adverse phenomena when considering hemin to be a viable treatment post MI.

## Author contributions

PM and SS contributed research conception and design. VA, RK, AF, EA and SS conducted experiments and acquired data. VA, RK and SS analysed data. VA, RK, PM and SS interpreted data. MC recruited patients and provided human samples. PM and SS secured funding and drafted the manuscript.

## Data availability statement

The data underlying this article are available in the article and in its online supplementary material.

