## Supplemental figures for "Regulation of BACH1 by hemin improves cardiac function in a mouse model of myocardial infarction"

**Supplementary methods**

*Cell transfection*

Opti-MEM media (ThermoFisher Scientific) and Lipofectamine RNAiMAX (Invitrogen) were used to transfect CP with On-target plus BACH1 SMARTpool siRNA (L-007750-00-0005, GE Healthcare), with Silencer® Select Negative Control No. 2 siRNA, (ThermoFisher Scientific) used as a control (final concentrations 25nM). The transfection reagent was removed after 6h and replaced with fresh media overnight. Media was then change to FBS-free media and cells incubated for 48h. RNA, protein and CM were then collected. Silencing was confirmed by qPCR and Western blotting.

*2-D Matrigel angiogenesis assay*

CAEC or CP were pre-treated with vehicle or hemin then seeded on the top of Matrigel® Growth Factor Reduced Basement Membrane Matrix (Corning) either in monoculture (4000 cells/well CAECs, 1500 cells/well CPs) or in coculture (4000 CAECs + 1500 PC/well), using Angiogenesis μ-Slides (IBIDI) and growth factors-free medium. Images were taken after 5 h using an inverted Leica microscope equipped with a 5× objective. The total tube length per imaging field was measured using ImageJ. To assess the interaction between PCs and CAECs, cells were labelled with green or red tracker Vybrant™ DiI Cell-Labeling Solution (Invitrogen; dilution 1:1000 in PBS, incubation for 5 min at 37°C followed by 15 min at 4°C).

**Supplementary figures**

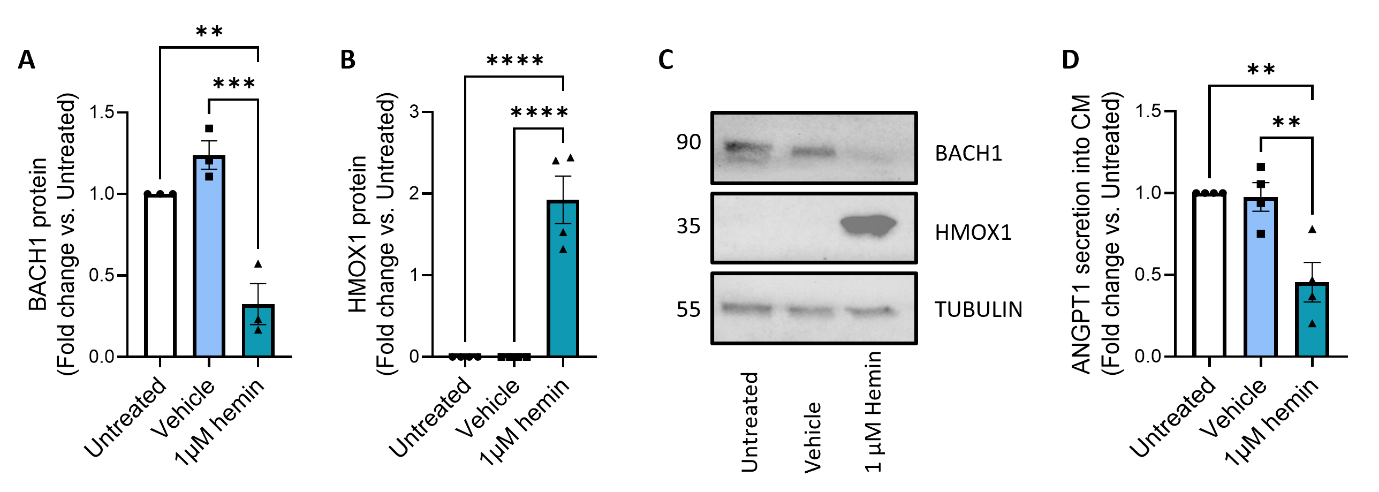

***Supplemental figure 1. Effect of 48h hemin treatment on BACH1 targets in CP.*** *Treatment of CP with hemin for 48h significantly decreased BACH1* ***(A)*** *protein expression and* ***(B)*** *significantly increased HMOX1 protein expression.* ***(C)*** *Representative Western blots.* ***(D)*** *ANGPT1 secretion in the conditioned media significantly decreased. One way ANOVA, ± SEM, n=3-4, **p<0.01 ***p<0.001 and ****p<0.0001 vs. cells only.*

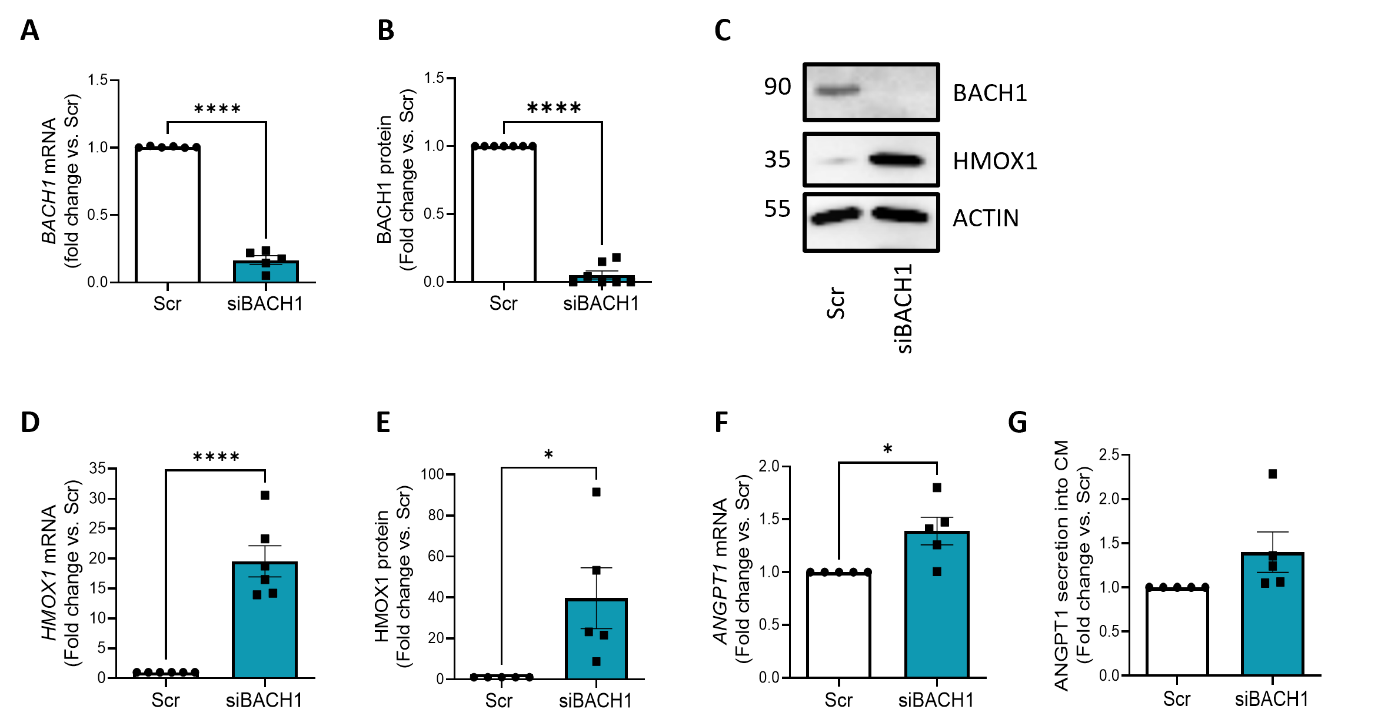

***Supplemental figure 2****.* ***Effect of BACH1 silencing using siRNA on expression of target factors in CP.*** *CP BACH1* ***(A)*** *gene and* ***(B)*** *protein expression significantly decreased in response to BACH1 silencing,* ***(C)*** *Representative Western blots.* ***(D)*** *HMOX1 gene and* ***(E)*** *protein expression significantly increased, as did ANGPT1* ***(F)*** *gene expression, however* ***(G)*** *there was no change in secretion of ANGPT1 into the conditioned media. All data unpaired t-tests, values ± SEM, n=3-6.  *P<0.05 and ****p<0.0001 vs. Scr.*

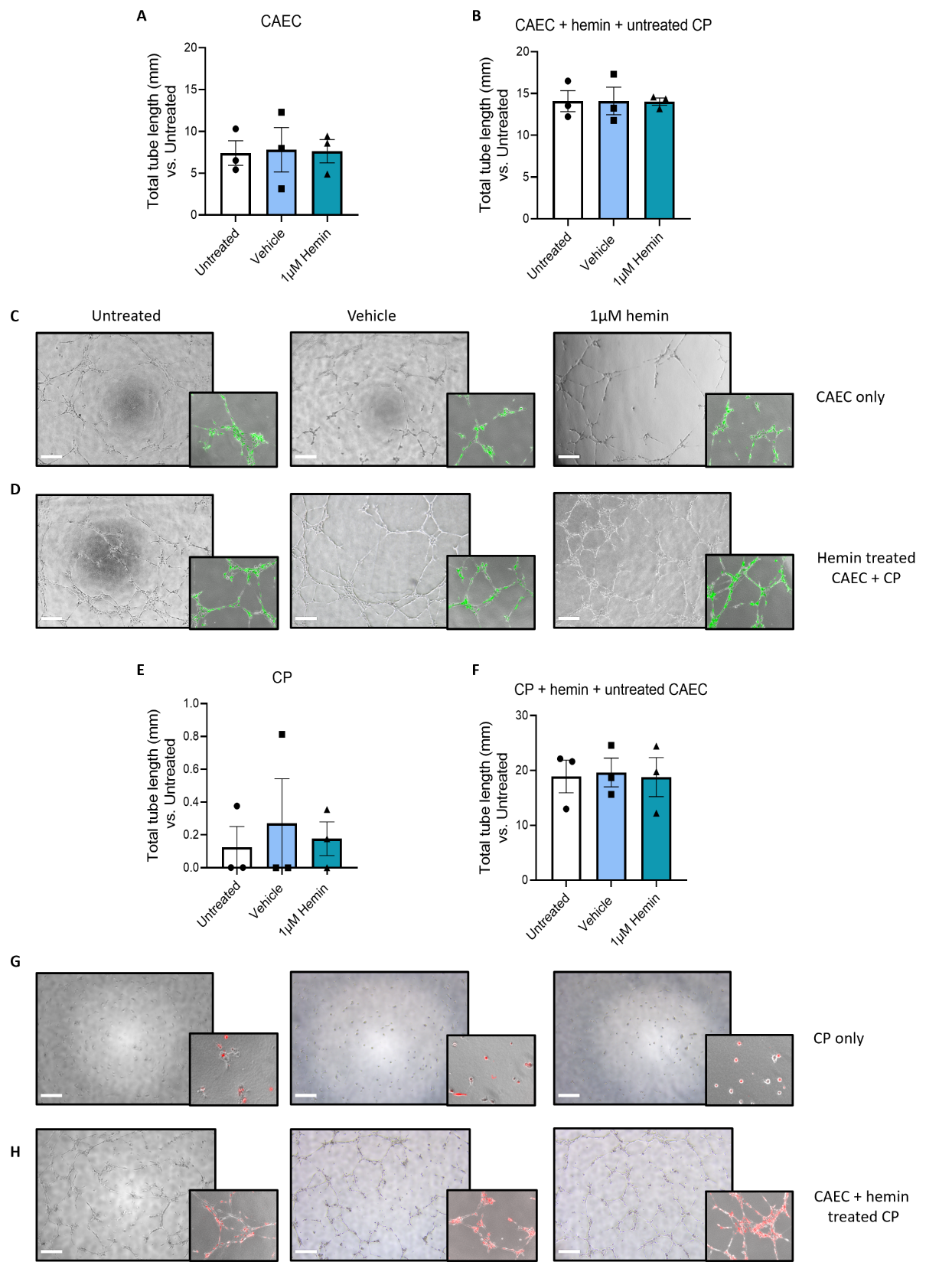

***Supplemental Figure 3. Effect of hemin treatment on vascular network formation.*** *Pre-treatment of CAEC with hemin for 24h prior to network formation experiment in* ***(A)*** *monoculture or* ***(B)*** *co-culture with untreated CP did not induce greater network formation. (****C&D, G&H)*** *Representative light microscopy and fluorescent images, green: CAEC, Red CP. Likewise pre-treatment of CP with hemin for 24h prior to network formation experiment in* ***(E)*** *monoculture or* ***(F)*** *co-culture with untreated CAEC did not induce greater network formation. One way ANOVA, ± SEM, n=3.*

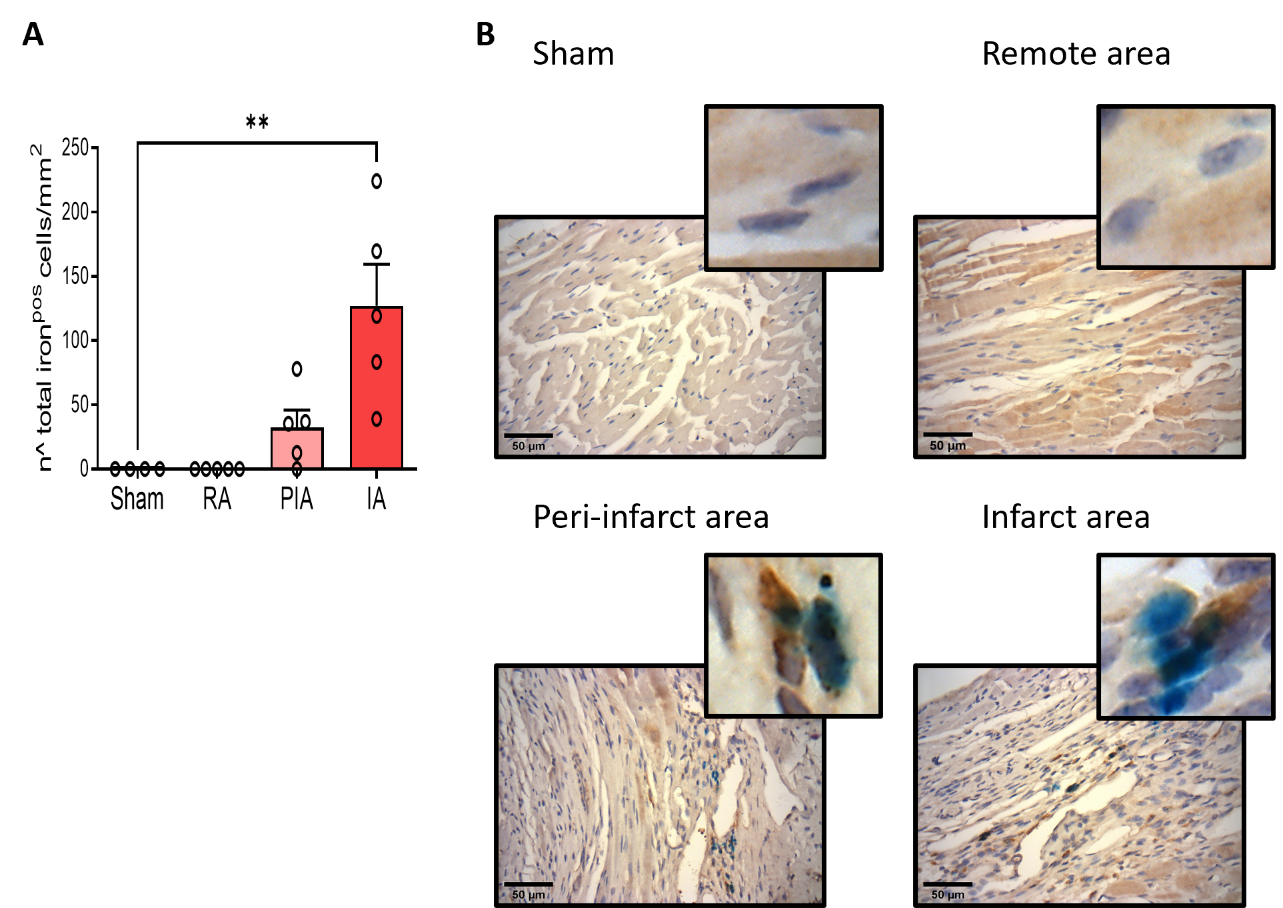

***Supplementary figure 4. Total iron accumulation in control sham and ischemic mouse samples. (A)*** *A significant increase of total iron accumulation was observed only in the infarcted area after MI.* ***(B)*** *Representative images. Blue: Iron, Brown: HMOX1, Purple: Haematoxylin nuclear stain. One-way Anova, ±SEM, n=5. **P<0.05*

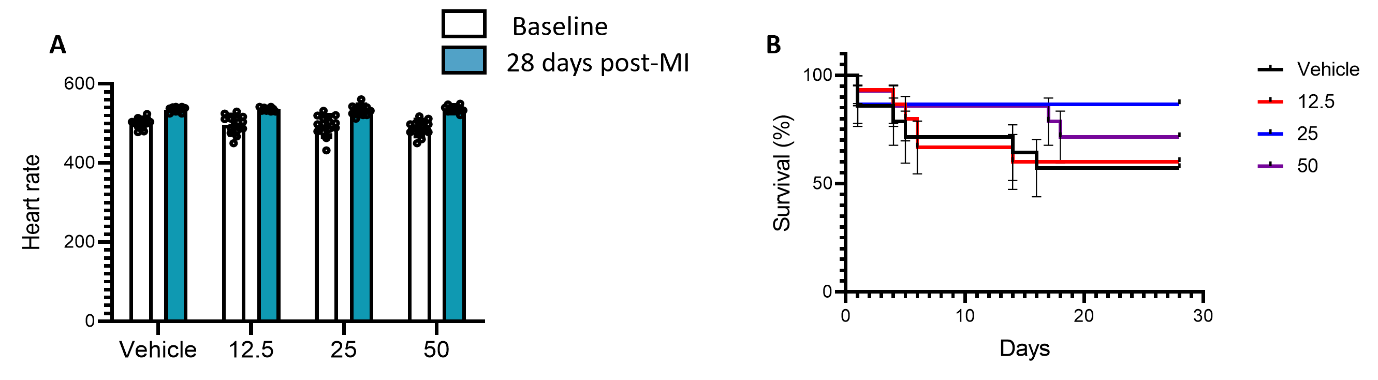

***Supplemental figure 5****.* ***Effect of Hemin treatment on the mouse heart.*** *There was no effect of treatment on* ***(A)*** *heart rate. Two way ANOVA, ± SEM, n=14. *P<0.05.* ***(B)*** *Mice treated with hemin also showed a trend for an improved survival rate, however Kaplan-Meier survival analysis showed there was no significant effect, ± SEM, n=14, vs. vehicle.*
